# The benefits and perils of import in small cattle breeding programs

**DOI:** 10.1101/2022.12.09.519737

**Authors:** J. Obšteter, J. Jenko, I. Pocrnic, G. Gorjanc

**Affiliations:** Department of Animal Science, Agricultural Institute of Slovenia, Ljubljana, Slovenia; Geno Breeding and A.I. Association, Hamar, Norway; The Roslin Institute and Royal (Dick) School of Veterinary Studies, University of Edinburgh, Edinburgh, United Kingdom; Biotechnical Faculty, University of Ljubljana, Ljubljana, Slovenia

**Keywords:** cattle breeding, import, genetic gain, partition analysis

## Abstract

Small breeding programs are limited in achieving competitive genetic gain and prone to high rates of inbreeding. Thus, they often import genetic material to increase genetic gain and to limit the loss of genetic variability. However, the benefit of import depends on the strength of genotype by environment interaction. It also also diminishes the relevance of domestic selection and the use of domestic breeding animals. Introduction of genomic selection has potentially execerbated this issue, but is also opening the potential for smaller breeding program.

The aim of this paper was to determine when and to what extent do small breeding programs benefit from import. We simulated two cattle breeding programs differing in selection parameters representing a large foreign and a small domestic breeding program that differ in the initial genetic mean and annual genetic gain. We evaluated a control scenario without the use of foreign sires in the domestic breeding program and 20 scenarios that varied the percentage of domestic dams mated with foreign sires, the genetic correlation between the breeding programs (0.8 or 0.9), and the time of implementing genomic selection in the domestic compared to the foreign breeding program (concurrently or with a 10-year delay). We compared the scenarios based on the genetic gain and genic standard deviation. Finally, we partitioned breeding values and genetic trends of the scenarios to quantify the contribution of domestic selection and import to the domestic genetic gain.

The simulation revealed that when both breeding programs implemented genomic selection simultaneously, the use of foreign sires increased domestic genetic gain only when genetic correlation was 0.9. In contrast, when the domestic breeding program implemented genomic selection with a 10-year delay, genetic correlation of 0.8 sufficed for a positive impact of import. In that scenario, domestic genetic gain increased with the increasing use of foreign sires but with a diminishing return. The partitioning analysis revealed that the contribution of import expectedly increased with the increased use of foreign sires. However, the increase did not depend on the genetic correlation and was not proportional to the increase in domestic genetic gain. This means that a small breeding program could be overly relying on import with diminishing returns for the genetic gain and marginal benefit for the genetic variability.

The benefit of import depends on an interplay of genetic correlation, extent of using foreign sires, and a breeding scheme. It is therefore crucial that small breeding programs assess the possible benefits of import beyond domestic selection. The benefit of import should be weighted against the perils of decreased use of domestic sires and decreased contribution and value of domestic selection.

## INTRODUCTION

Cattle breeding programs import genetic material to increase genetic gain and genetic diversity. Small breeding programs depend more on import since they usually have lower rates of genetic gain due to lower intensity and accuracy of selection, and slower technological and methodological development. They might also have higher rates of inbreeding due to a smaller genetic pool. The benefit of import depends on the difference in genetic means, difference in breeding objectives, and the presence of the genotype-environment interaction (GxE) between the foreign and domestic populations. GxE can reduce the benefit of the import and downstream selection of imported germplasm due to different genetic value of the germplase in different environments (Falconer and Mackay, 1996). We often measure GxE with genetic correlation of a trait in different environments. The correlation of less than 1 implies the presence of GxE, which can cause a re-ranking of genotypes in different environments. If environments differ sufficiently, import may not be beneficial. To maximise the short-term genetic gain in breeding programs of equal size with conventional selection, the correlation between environments needs to be at least 0.7 – 0.8 (Banos and Smith, 1991; Robertson, 1959), and more than 0.9 to allow for a long-term cooperation between breeding programs (Cao et al., 2020; Mulder and Bijma, 2006). This means that if the correlation is less than 0.9, populations should stop importing after a couple of generations and rely entirely on their own genetic improvement. Recent introduction of genomic selection enabled the long-term cooperation at slightly lower correlations, between 0.85 and 0.875 for populations of equal size (Cao et al., 2020). When cooperating populations are of unequal size, the smaller population with a lower genetic mean benefits more from the exchange of genetic material than the larger population (Smith and Banos, 1991; Mulder and Bijma, 2006), particularly with genomic selection (Cao et al., 2020).

The import benefits the genetic gain and genetic diversity of small breeding programs, but also diminishes the importance of domestic breeding activities, if not even whole breeding programs. The benefits of import therefore need to be weighted against the eventual loss of the domestic breeding programs, which are difficult to quantify. Gorjanc et al. (2011, 2012) analysed the contribution of different countries to international and national genetic trends in Brown Swiss and Holstein breeds. They showed a major contribution of breeding activities originating in the USA despite a multitude of national breeding programs, each running its own domestic selection in addition to import. The missed opportunity of domestic selection was also shown in Fetherstone et al. (2021). They inspected the genetic and economic benefit of foreign sires to the domestic sheep industry with the gene-flow method. They showed that using foreign genetics resulted in a lower benefit than increasing the use of progressive domestic selection. However, genomic selection offered new opportunities for cooperation between populations in two ways. First, shorter generation intervals with genomic selection allow for a faster dissemination of imported genetic material in breeding and commercial parts of domestic population. And second, breeding programs can exchange summary statistics obtained from genome wide association studies, which can increase accuracy of selection (Vandenplas et al., 2018) and to potentially manage GxE (Mulder, 2016). Such an exchange of genomic or association information alone can increase genetic gain even when genetic correlation is as low as 0.4 (Slagboom et al., 2019).

The aim of this paper was to determine when and to what extent do small cattle breeding programs benefit from import. To this end this paper: i) tests the effect of increased use of foreign sires on genetic gain and genetic variability in the domestic population at different genetic correlation between populations; and ii) partitions genetic trends of the tested scenarios to quantify the contributions of domestic and foreign populations to the domestic genetic gain.

## MATERIALS AND METHODS

The study aimed to quantify the effect of varying use of foreign sires on genetic gain and genetic variability in a small cattle breeding program. To this end we simulated two breeding programs differing in selection intensity and accuracy to mimic the existence of a large foreign and a small domestic breeding program that differ in the initial genetic mean and annual genetic gain. We evaluated 20 scenarios in which the domestic breeding program used foreign sires. The scenarios differed in (i) the percentage of domestic females mated with foreign sires, (ii) the genetic correlation between the domestic and foreign breeding program, and (iii) the delay in implementing genomic selection in the domestic compared to the foreign breeding program. We compared the scenarios in terms of genetic gain and genetic variability in the domestic population. Finally, we partitioned the trends of genetic gain of the tested scenarios to quantify the contributions of domestic and foreign selection.

### Simulation of the base population and historical breeding

We simulated two populations with overlapping generations that mimicked a domestic and a foreign breeding program and differed in selection intensity and accuracy. The simulation included genomes, genetic values, and phenotypes for a polygenic trait with heritability 0.25, and a whole cattle breeding program as described previously (Obšteter et al., 2019). In short, both breeding programs started from the same base population. For computational simplicity, each breeding program included ∼22,500 active animals but differed in the selection parameters to reflect their size and resources. The simulated trait represented a polygenic net merit encompassing many traits influenced by 10,000 causal loci (Obšteter et al., 2019). To generate genetic difference between the two breeding programs due to genetic drift and selection, we ran each breeding program for 20 years of random mating followed by a 10 year burn-in period with conventional selection with progeny testing in both populations. We simulated populations with overlapping generations. In both population, we generated 4320 females calves every year, culled 11.2% due to problems at birth, culled 20% of cows after each lactation, and culled all the remaining cows after the forth lactation. In the domestic breeding program, we yearly selected 139 bull dams out of cows in the second, third, and fourth lactation. We then generated 45 male calves from matings of bull dams and progeny tested sires (elite matings). Out of these we chose 8 for progeny testing of which 4 were eventually selected as sires for widespread insemination of cows. In the foreign population, we implemented a higher selection intensity and accuracy to achieve a difference of 1.5 genetic standard deviation in the initial genetic means between the two populations. To this end, we yearly selected 855 bull dams to produce 300 offspring of elite matings, of which we selected 150 for progeny testing, and 20 as elite sires. All elite sires in the domestic and foreign population were kept in use for 5 years. In the period of historical breeding, all selections in the domestic breeding program were done on pedigree-based estimates of breeding values, and all selections in the. foreign breeding program were done on true breeding values.

We simulated a genetic correlation between the two breeding programs to be either 0.8 or 0.9. We simulated the genetic correlation as two genetically correlated traits that represented the net merit trait expressed in the domestic (“domesticTrait”) and foreign (“foreignTrait”) breeding program. AlphaSimR simulates the genetic correlation by correlating the QTL effects for the traits. We selected animals in the domestic breeding programs based on the trait expressed in the domestic environment, and animals in the foreign breeding programs based on the genetically correlated trait expressed in the foreign environment.

### Scenarios

We next simulated an additional 20 years in which we evaluated 24 different scenarios of importing foreign sires into the domestic breeding program, including a control scenario without import. The scenarios differed in three ways. First, they varied the genetic correlation as explained above (r_g_ of and 0.9) between the net merit trait expressed in the two breeding programs. Second, the scenarios varied the use of foreign sires in the domestic breeding program. We mated 0%, 10%, 25%, 50%, or 100% of the domestic females with foreign sires (from here on we use the term % foreign sires). We additionally created a scenario that mated only elite bull dams with foreign sires. Third, the scenarios varied in introducing genomic selection, where (i) both breeding programs implemented genomic selection simultaneously in year 30; and (ii) the foreign breeding program implemented genomic selection in year 30, while the domestic breeding program implemented it with a 10 year delay, in year 40. The latter mimicked a situation where small breeding programs adopt a new technology with a delay due to the lack of resources, technical support, and scepticism. With genomic selection in the domestic breeding program, we genotyped all offspring of elite matings (45) and selected the top 6 as elite sires. In the foreign breeding program, we increased the number of offspring from elite matings to 400, genotyped them all, and selected the top 15 as elite sires. This scheme mimicked the fact that large breeding programs usually take a better advantage of the decreased cost of genomic compared to conventional testing and increase the intensity of sire selection in genomic selection (Wiggans et al., 2017). We replicated the simulation of base population followed by historical breeding and each scenario ten times. Construction of training populations and genetic evaluation with genomic data is described in the following. All selections were done on either pedigree-or genomics-based estimates of breeding values for the net merit for the corresponding trait.

### Estimation of breeding values

We estimated breeding values either with the pedigree or genomic single-step genetic evaluation. Breeding values were estimated within each breeding program without sharing data and using all available pedigree and phenotypic data, and a collection of genomic data. We used blupf90 (Misztal et al., 2018) with default settings for the estimation of breeding values in each generation of selection and variance components used for the simulation of the traits. Initial set of genomic data consisted of genotypes from all active cows (∼10,000 in each population) and all elite sires (80 in the small and 200 in the large population). We updated the genomic data every year by removing genotypes from the oldest generation of cows and replacing them with genotypes from the new first lactation cows. We also added genotypes from every new generation of male candidates.

### Analysis of scenarios

We compared the scenarios based on genetic gain, loss of genetic variability, and contribution of domestic selection and import to genetic gain. We computed the genetic gain in each breeding program by averaging true breeding values by the birth year, and standardized it to have the mean of 0 and standard deviation of 1 in the domestic breeding program in the first year of comparison (year 30). We standardized the genetic gain in the foreign breeding program with the values from the domestic breeding program to facilitate comparison. We expressed annual genetic gain by regressing the average true breeding value onto the birth year of animals. We computed genic variance as a measure of genetic variablity as 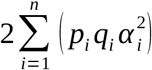 where n is the number of causal loc, *p* and *q* are the allele frequencies, and α is the allele substitution effect at the causal loci. We standardized the genic variance to have the value of 1 in the domestic breeding program in the first year of comparison. We used AlphaPart R-package (Obšteter et al., 2021) to partition the trend in genetic gain of the domestic breeding program into the contribution of domestic selecton and import from the foreign breeding program. We partitioned the true breeding values and hence the true genetic trend to evaluate the impact of import without the uncertainty and any bias in estimated breeding values. To obtain genetic contributions (i.e. proportion of domestic and foreign genes) to the domestic population, we partitioned a vector of ones.

## RESULTS

When both populations implemented genomic selection simultaneously, using foreign sires significantly increased domestic genetic gain only when genetic correlation between populations was 0.9. When the domestic population implemented genomic selection 10 years after the foreign population, using foreign sires significantly benefited the domestic genetic gain at both tested genetic correlations, 0.8 and 0.9. When the use of foreign sires was beneficial, the domestic genetic gain increased with increasing use of foreign sires but with a diminishing return. The use of foreign sires also marginally alleviated the loss of genic variance in the domestic population at both tested genetic correlations. The partitioning analysis revealed that the contribution of foreign population to domestic genetic gain increased with increasing use of foreign sires. However, the increase in foreign contribution was not proportional to the increase in genetic gain.

We first compare the scenarios in terms of the genetic gain and genic standard deviation in the last year of simulation, year 50, as well as the annual genetic gain when appropriate. For scenarios that changed the domestic breeding scheme in year 40 we also report the results for year 40. We next compare the scenarios in terms of partitioned trend in genetic gains.

### Genetic gain and genetic variability

#### Simultaneous implementation of genomic selection in both populations

Import has increased the domestic genetic gain when genetic correlation with the foreign population was 0.9 or more (Figure 1). When genetic correlation was 0.8, import did not significantly increase domestic genetic gain compared to only using domestic genomic selection. When genetic correlation was 0.9, the domestic genetic gain increased with increasing import, but with diminishing returns (Figure 1 and Table S1). Compared to only domestic selection, genetic gain significantly increased by 10% (measured at year 50) when we mated 25% dams with foreign sires. Increasing this percentage to 50% and 100% respectively increased genetic gain to 16% and 18%. Mating only bull dams with foreign sires gave a domestic genetic gain similar to when mating 50% dams with foreign sires. Although the differences were small, results suggest that using a mix of foreign and domestic sires could alleviate the loss of genic standard deviation in the domestic population (Figure 2 and Table S2). At both genetic correlations, we observed the largest loss of genic standard deviation by year 50 in the scenario that did not import foreign sires (3-4%), and the smallest loss in the scenarios that mated bull dams exclusively with foreign sires (1-2%).

**Figure 1:**
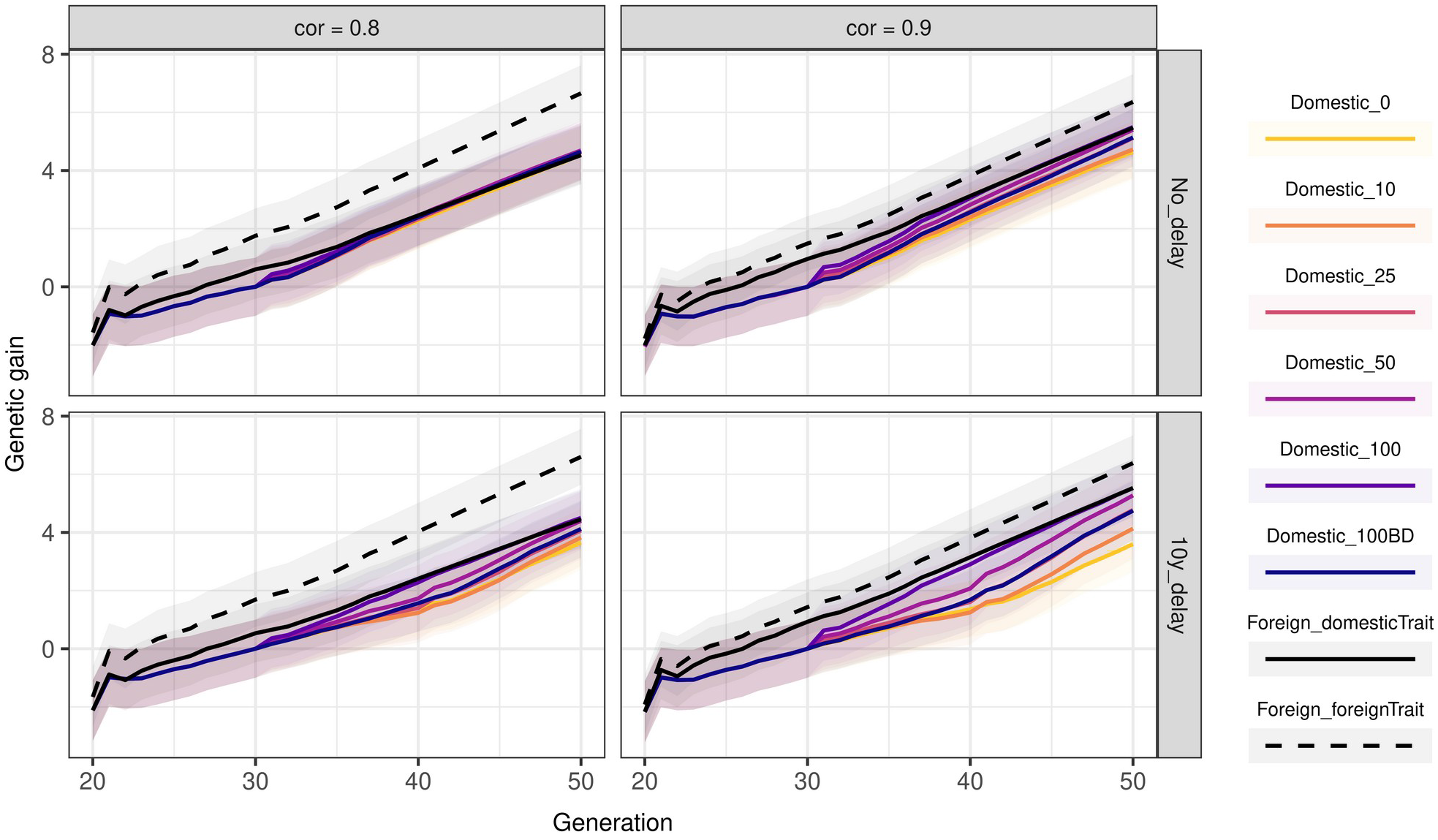
Genetic gain by scenario, genetic correlation between populations, and the time of implementing genomic selection. The lines represent the average breeding value by the year of birth and the ribbon the corresponding standard deviation across 10 replicates. The lines named “Domestic” show the domestic genetic gain with varying percentage of domestic females mated with foreign sires (0, 10, 25, 50, 100). Scenario 100BD mated only the elite bull dams with foreign sires. The lines named “Foreign” show the genetic gain of the foreign population for the trait expressed on the scale of the domestic (“domesticTrait”) and the foreign (“foreignTrait”) breeding program. Scenarios marked with “No_delay” implemented genomic selection in both populations simultaneously (in year 30). Scenarios marked with “10y_delay” implemented genomic selection in the domestic population 10 years after the foreign population (in year 40).

**Figure 2:**
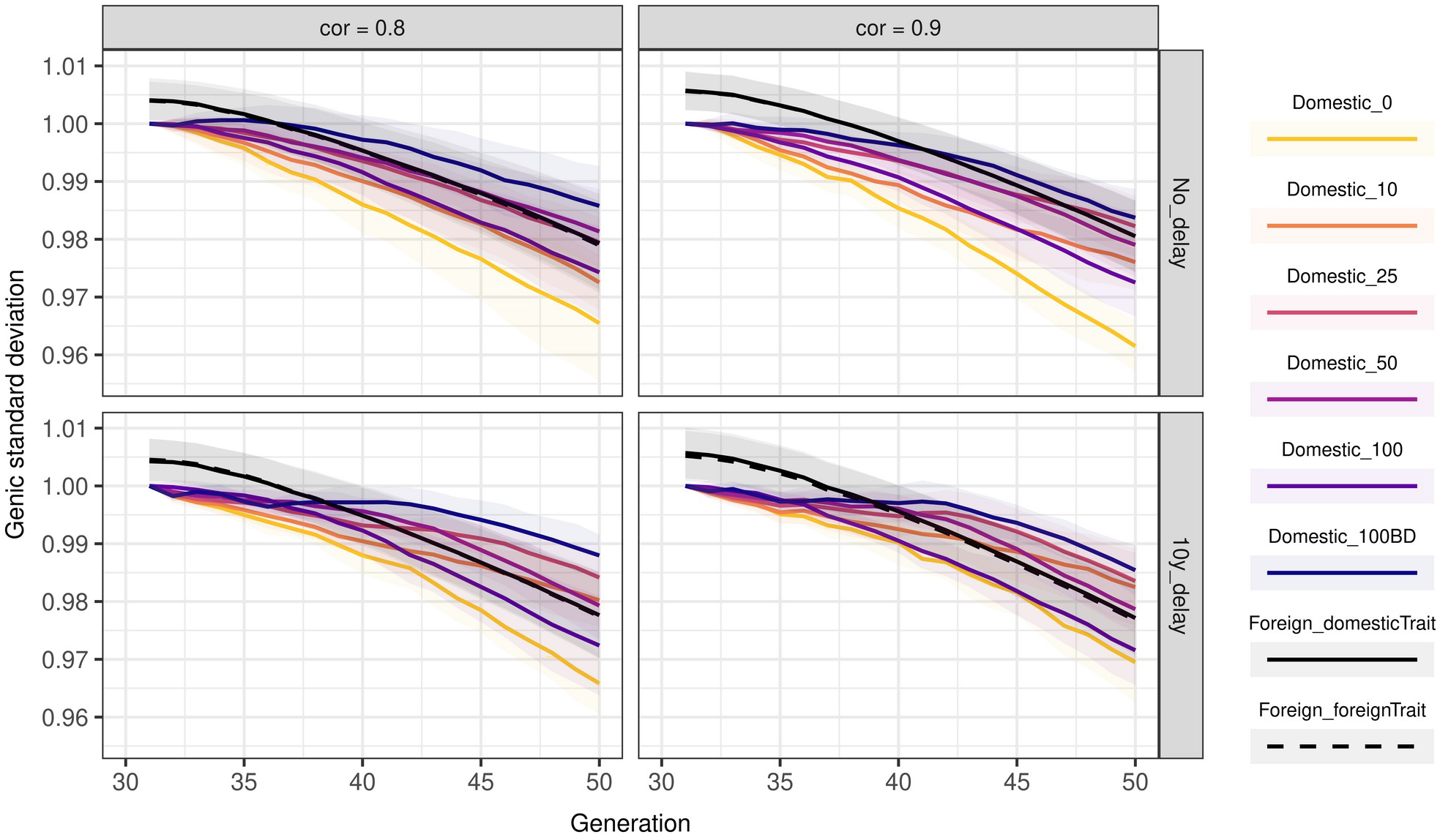
Genic standard deviation by scenario, genetic correlation between populations, and the time of implementing genomic selection. The lines represent the average genic standard deviation by the year of birth and the ribbon the corresponding standard deviation across 10 replicates. The lines named “Domestic” show the domestic genic standard deviation with varying percentage of domestic females mated with foreign sires (0, 10, 25, 50, 100). Scenario 100BD mated only the elite bull dams with foreign sires. The lines named “Foreign” shows the genic standard deviation of the foreign population for the trait expressed on the scale of the domestic (“domesticTrait”) and the foreign (“foreignTrait”) breeding program. Scenarios marked with “No_delay” implemented genomic selection in both populations simultaneously (in year 30). Scenarios marked with “10y_delay” implemented genomic selection in the domestic population 10 years after the foreign population (in year 40).

#### 10-year delay in the implementation of domestic genomic selection

When the domestic population implemented genomic selection 10 years after the foreign population, using foreign sires increased the domestic genetic gain at both tested genetic correlations (Figure 1 and Table S3). The increase was however larger when the genetic correlation with the foreign population was 0.9. The domestic genetic gain increased with increasing the use of the foreign sires.

Implementing domestic genomic selection changed the effect of using foreign sires and hence we compare the scenarios based on the genetic gain in years 40 and 50 for period prior and after the implementation, respectively (Figure 1 and Table S3). By year 40, import increased the domestic genetic gain between -12% and 62% when genetic correlation was 0.8 and between -8% and 114% when genetic correlation was 0.9 compared to the scenario without the import. During the years 40-50 the difference between the scenarios remained more constant due to less variable annual genetic gains (Figure S1). By year 50, import increased the domestic genetic gain between 5% and 23% when genetic correlation was 0.8 and between 15% and 53% when genetic correlation was 0.9 compared to the scenario without the import. The increase was significant when we used at least 10% or 25% of foreign sires respectively with genetic correlation 0.8 and 0.9 (Figure 1, Table S3). Mating only bull dams with foreign sires resulted in the genetic gain comparable to when we used 25% of foreign sires.

As in the scenarios that implemented the genomic selection simultaneously in both populations, the loss in genic standard deviation was small in all scenarios (Figure 2 and Table S4). Again, the results suggest that using a mix of domestic and foreign sires can reduce the loss of genetic variability. We observed the largest loss by year 50 (3%) in the scenario that did not use foreign sires and in the scenario that used exclusively foreign sires. This was observed at both tested genetic correlations. When we mated only bull dams with foreign sires the loss was only 1%.

### Contribution of domestic and foreign selection

The contribution of domestic selection and import to the domestic genetic trend is shown in figures 3 and 4. The proportion of domestic and foreign genes in the domestic population is shown in figures S4 and S5. The contribution of import to domestic genetic gain, as well as the proportion of foreign genes, expectedly increased with increasing import. This was so regardless of the genetic correlation between the populations or the breeding scheme.

**Figure 3:**
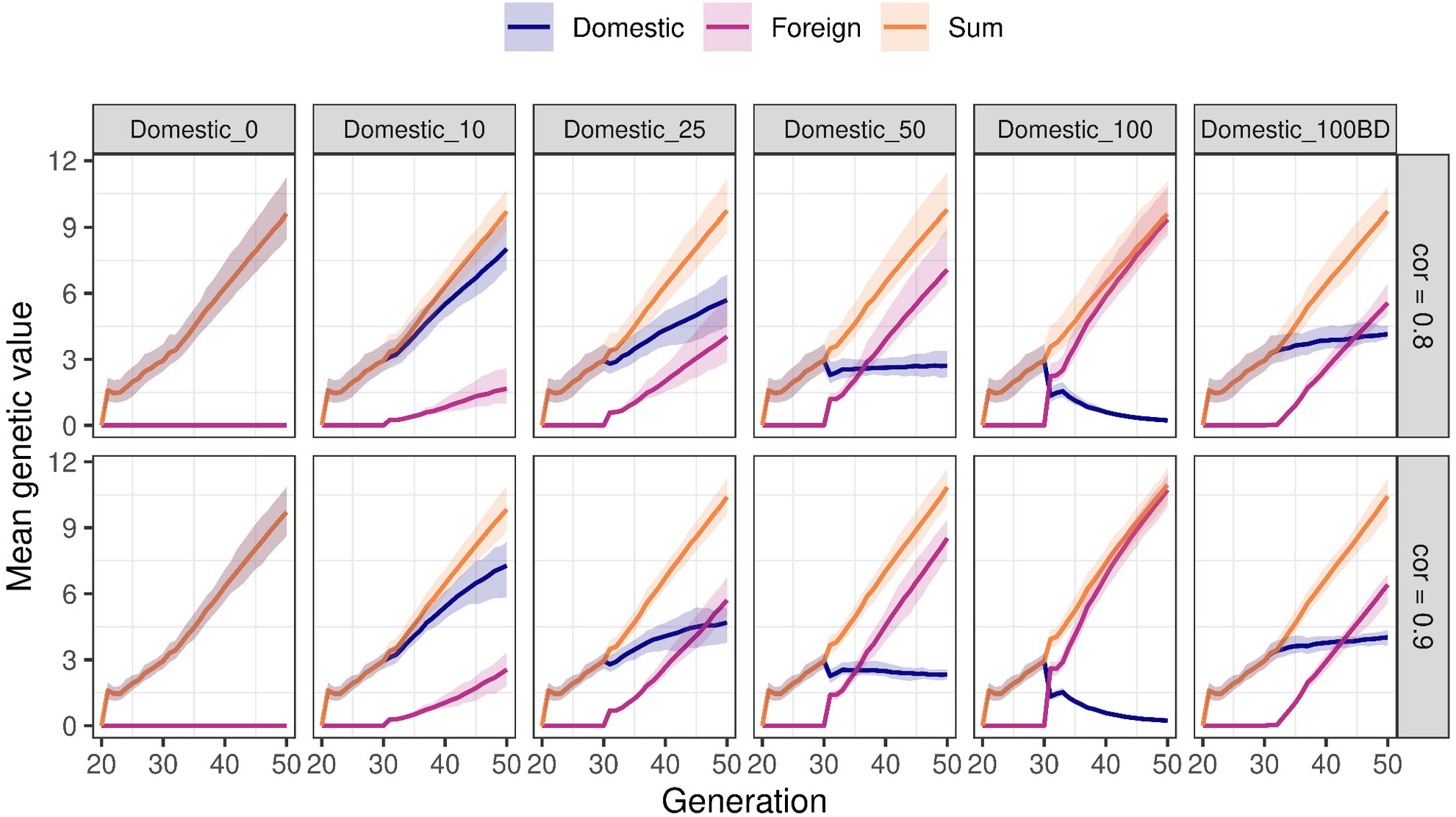
Partitioning of the domestic genetic trend into domestic and foreign contributions in scenarios with simultaneous implementation of genomic selection in both populations. The scenarios first varied the percentage of domestic females mated with foreign sires (0, 10, 25, 50, or 100). In 100BD scenario we mated only bull dams with foreign sires. Second, the scenarios varied genetic correlation between two breeding programs (0.8 or 0.9).

**Figure 4:**
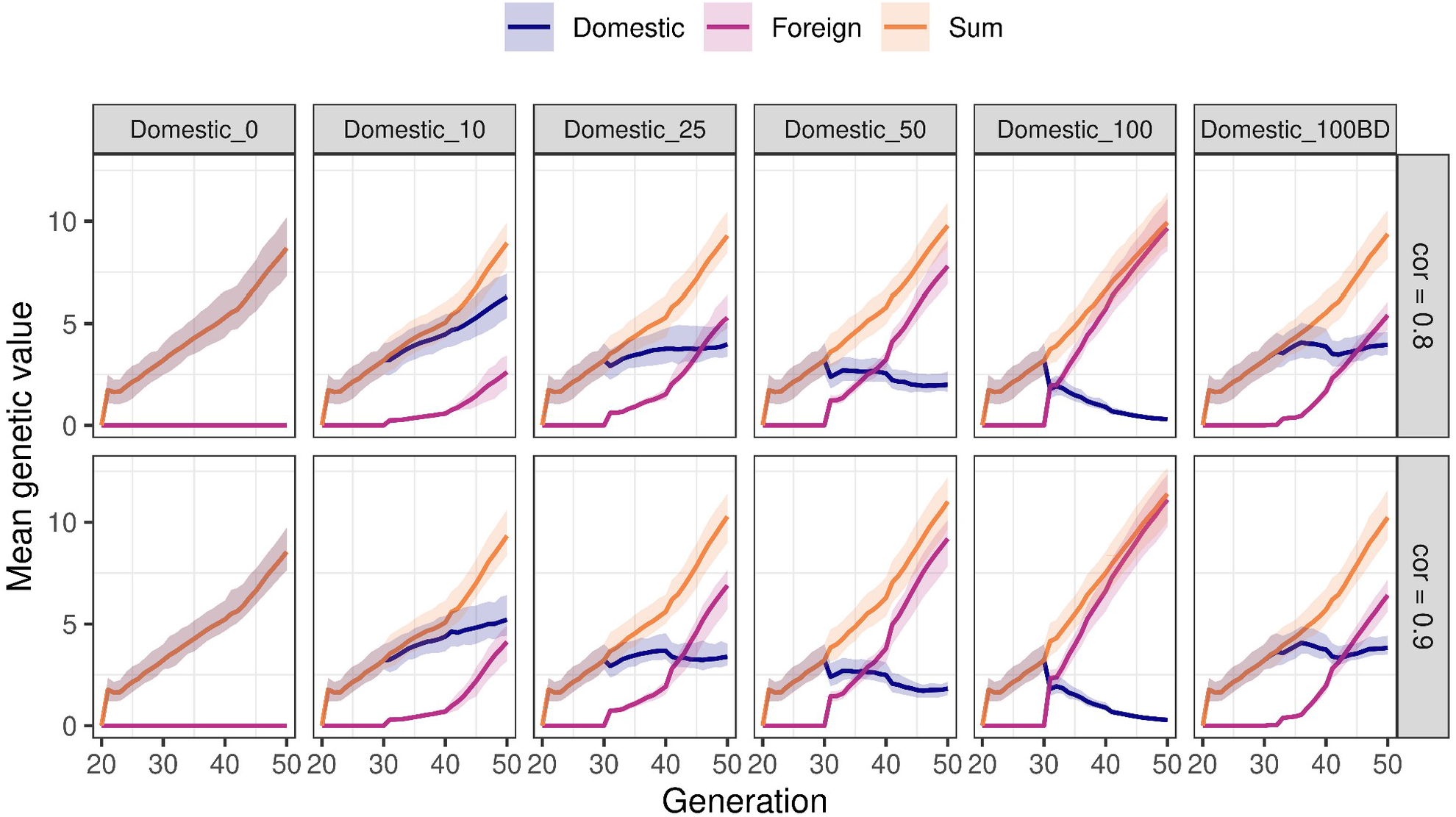
Partitioning of the domestic genetic trend into domestic and foreign selection in scenarios with a 10-year delay in the implementation of genomic selection in the domestic population. The scenarios first varied the percentage of females mated with foreign sires (0, 10, 25, 50, or 100). In 100BD scenario we mated only bull dams with foreign sires. Second, the scenarios varied genetic correlation between two breeding programs (0.8 and 0.9). Foreign population implemented genomic selection in year 30 and domestic population in year 40.

#### Simultaneous implementation of genomic selection in both populations

The partitioning analysis revealed that the contribution of import to the domestic genetic gain expectedly increased with the increasing import (Figure 3). In contrast to genetic gain, the contribution of import to domestic genetic gain increased regardless of the genetic correlation, although it was higher and increased faster when genetic correlation was 0.9 (Figure 3 and Figure S2). In year 50, across the scenarios foreign population contributed between 17% and 98% to the domestic genetic gain when genetic correlation was 0.8, and between 26% and 98% when the genetic correlation was 0.9.

The scenario that used 25% of foreign sires represented a breaking-point. When we used less than 25% of foreign sires, the contribution of import increased at a slower rate than the contribution of domestic selection (Figure S2). When we used 25% or more foreign sires, the contribution of import started to increase at a faster rate than the domestic contribution, which means that it eventually contributed majority to the domestic genetic gain. Mating 50% of domestic females with foreign sires represented a point at which the domestic contribution was practically constant throughout the years (Figure 3, Figure S2) and all the gain was contributed by the import. When we mated all the domestic females with foreign sires, the domestic contribution decreased since we did not use domestic sires for breeding. The scenario that mated only the bull dams with foreign sires had comparable foreign contribution as the scenarios that 25% of the domestic females with foreign sires.

#### 10-year delay in the implementation of domestic genomic selection

As in the scenarios with the simultaneous implementation of genomic selection, increased import expectedly increased the contribution of foreign population regardless of the genetic correlation between the populations (Figure 4 and, Figure S5). Across scenarios, the foreign contribution to the domestic genetic gain in year 50 was between 29% and 97% when genetic correlation was 0.8, and between 44% and 98% when genetic correlation was 0.9.

The dynamics of the domestic and foreign contributions differed in years 30-40 (domestic conventional selection) compared to years 40-50 (domestic genomic selection). In years 30-40, we had to use 25% of foreign sires for the foreign contributions to increase faster than the domestic (Figure 4 and Figure S3). In years 40-50, the foreign contribution increased faster than domestic already at 10% of foreign sire use. A higher rate of increase of foreign contribution resulted in the foreign population eventually contributing the majority to the domestic genetic gain.

## DISCUSSION

Our results show that the benefit of using foreign sires in small cattle populations depends on the genetic correlation between the populations, the extent of the use of foreign sires, and the breeding scheme in each of the populations. Small domestic populations can be limited in achieving genetic gain due to limited intensity and accuracy of selection, and can be also more prone to high rates of inbreeding. Thus, they often use import to increase genetic gain and genetic variability. However, import can diminish the importance of the domestic breeding program. It is therefore crucial that small populations quantify the benefit and the realized contribution of import to the domestic genetic gain. Our results raise three discussion points: 1) how does varying the genetic correlation between populations affect the domestic genetic trend and the foreign contribution; 2) how does varying the use of foreign sires affect the domestic genetic trend and the foreign contribution; and 3) how does late implementation of genomic selection in the domestic population affect the domestic genetic trend and the foreign contribution.

### Genetic correlation

The benefit of using foreign sires to increase the domestic genetic gain increased with increasing genetic correlation. In some settings with genetic correlation of 0.8, the use of foreign sires even resulted in zero benefit. This shows the importance of assessing the genetic correlation with potential foreign suppliers of genetic material before import. Slagboom et al. (2019) showed that when populations are of different size, having a joint breeding program is only beneficial for the small breeding program when the genetic correlation is above 0.79. Other studies suggested a lower correlation is sufficient for populations of unequal size (Cao et al., 2020; Mulder and Bijma, 2006; Robertson, 1959). They suggested a genetic correlation between 0.6 and 0.7 suffices for short-term cooperation and between 0.7 and 0.8 for long-term cooperation, which was not confirmed by our results. Our results agree with Schmidtmann et al. (2021), who explored the exchange of sires between Red dairy and dual-purpose breeds. They first showed that the benefit of cooperation strongly depends on the trait and weighing of the selection indices. Second, they showed that unless the correlation is close to unity, the populations will in long-term rely completely on their own breeding program, despite initial exchange and benefit. Vargas and Arendonk (2004) also showed that when the genetic correlation is 0.75, importing semen from a larger country with a higher genetic mean achieves the same genetic response as performing progeny testing with smaller progeny groups in the domestic population, which is in line with our results, but the extent of import is important as discussed below. Many breeding programs in the Global North have high genetic correlations in range between 0.85 and 0.90 (Fikse et al., 2003) There are some exceptions with correlations below 0.8 (Rekaya et al., 2001, Harris, 2005; Interbull, 2021). The environment and associated GxE is of particular importance when considering exchange of genetic material between the Global North and the Global South. For example, the genetic correlation between Kenya and UK was found to be only 0.49 (Ojango and Pollott, 2002).

Even though in our study genetic correlation of 0.8 diminished or even eliminated the benefit of import for the domestic genetic gain, the partitioning analysis revealed that the foreign contribution to domestic genetic gain still increased and so did the percentage of foreign genes in the domestic population. This shows that the foreign sires were selected based on the superiority of their breeding values for the trait in the foreign population that did not translate into superiority for the trait in the domestic population (Figure S6). The increase in the foreign contribution was driven by the import of sires directly, but also by the domestic sires being offspring of foreign sires. Results showed that when we imported 50% of foreign sires, additional ∼50% of the domestic sires were offspring of the foreign sires (same in other scenarios with the corresponding percentage). Percentage of domestic sires sired by foreign sires was slightly higher when genetic correlation was 0.9 than when genetic correlation was 0.8. Same was observed at other import percentages. This has important implications for breeding programs in small populations since they could be importing genetic material and genetically converting the domestic population into the foreign population without benefits for the genetic gain and marginal benefit for the genetic variability. Increasing the foreign contribution to the domestic genetic gain diminishes the importance of domestic breeding and decreases the adaptation of the population to the local environmental conditions. Schmidtmann et al. (2021), also showed that intensive imports lead to genetic convergence of populations, which reduces global genetic diversity.

Important novelty in this study is the analysis of contribution of different groups (domestic and foreign) to the genetic gain. Although previous studies did not perform this analysis, they still analysed the optimal percentage of import. Banos and Smith (1991) explored the use of sires in two populations of equal size and equal initial genetic mean with selection across populations. They showed that when the genetic correlation is 0.8, the populations gradually use more of their own bulls and rely almost completely on domestic selection by generation 5. When genetic correlation is the switch to using only domestic bulls is slower and the populations use about 75% of their own bulls by generation 5. Slagboom et al. (2019) showed that with two environments and allowing for selection across them, the environments only started to select bulls across environments when genetic correlation was above 0.8. Similarly, in Slagboom et al. (2021) they showed that when two environments of equal size account for the GxE and select each on its own selection index, no bulls are selected across environments until genetic correlation was 0.7. Schmidtmann et al. (2021), also showed that only when the correlation between the population with different breeding goals is close to unity, they would colaborate in long-term with 30% import. This is not in concordance with our results showing that the foreign contribution increases regardless of the genetic correlation. In our case, we considered populations of unequal breeding nucleus size and different genetic means, which could increase the benefit of using foreign sires. Although we did not implement selection across populations or allow for a population to choose or modify the use of foreign sires, the domestic breeding program was free to choose the offspring of foreign or domestic sires as the new generation of domestic selection candidates.

The import also decreased the loss of genetic variability, although the loss was small in all tested scenarios. However, the genetic correlation did not significantly affect the loss of genetic in the correposnding scenarios.

### Extent of the foreign sire use

Increasing the use of the foreign sires in the domestic population increased the domestic genetic gain. In most scenarios, we needed to inseminate at least 25% cows with foreign sires to significantly increase genetic gain (Table S1). However, increasing the use of foreign sires beyond 25% or 50% was not beneficial in most of the scenarios. This is partly in agreement with Cao et al. (2020) who showed that when genetic correlation allows for a long-term cooperation between populations of unequal size yet equal genetic means, the optimum use of foreign sires in the smaller population is between 30% and 70% in conventional selection and between 60% and 80% in genomic selection. This means that the same genetic gain can be achieved with buying less foreign semen and investing funds into other breeding activities. Similarly, inseminating only bull dams with foreign sires has achieved the same genetic gain as mating 25% of all cows with the foreign sires in terms of genetic gain. In our simulation, this scenario required buying foreign semen for only 90 successful pregnancies versus ∼2,500 successful pregnancies with 25% cows.

The partitioning of the genetic trend expectedly showed that the contribution of import to domestic genetic gain significantly increases with the increasing use of foreign sires, but the genetic gain does not increase correspondingly. For example, using 10% of foreign sires did not significantly increase genetic gain in most of the settings, but it increased the foreign contribution to domestic genetic gain between 17% to 44%. The reason for this is that in addition to the 10% of directly imported sires, additional up to ∼10% of the domestic candidates were offspring of these imported bulls. For the same reason, increasing the use of foreign sires from 50% to 100% did not significantly increase genetic gain, but increased the foreign contribution by additional 20% to 32%. The main peril therefore is that breeding programs could be importing genetic material without any benefits and decreasing the importance of domestic breeding efforts for the local environment as already discussed above (in Genetic correlation).

Using 25% of foreign sires proved suitable for increasing domestic genetic gain while maintaining the contribution of domestic selection. We achieved the same result by mating only bull dams with foreign sires. This partly agrees with Banos and Smith (1991) who showed that populations with cooperative selection use between 25% and 30% of the sires from the other populations. But since they only evaluated five generations the percentage could decrease below that in the following generations.

The results also show that import reduces the loss of genic variance, although the loss was small in all the scenarios. The scenarios that used either 25% of foreign sires or used them only for bull dams maintained the most genic variance in all the settings. In contrast, in most settings using exclusively foreign fires (100% import) did not significantly increase the genic variance compared to the scenario that did not use foreign sires.

### Implementation of genomic selection in domestic population

We showed that the late implementation of genomic selection in the domestic population changes the effect of genetic correlation and extent of foreign sire use on domestic genetic gain. When the domestic population implemented genomic selection 10 years after the foreign population, the genetic correlation of 0.8 sufficed for the domestic population to benefit from the use of foreign sires. The first reason for this is that progeny-testing results in a lower genetic gain compared to genomic breeding scheme due to a longer generation interval, which increases the benefit of using foreign sires despite lower genetic correlation. The second reason is the decreased accuracy of the progeny testing in the domestic population. Since we simulated a population of a constant size, the use of foreign sires decreased the number of domestic females available for mating with domestic sires and the size of the progeny groups for progeny testing.

The domestic annual increase in genetic gain with genomic selection differed depending on whether it was preceded by 10 years of conventional selection with importing or not (Figure 1, Figure S1). This could be explained by the following. When we ran domestic conventional selection in years 30-40, we imported sires that were selected due to the superiority of their genomic segments, that is their genomic breeding value in the foreign population. But in the domestic population, we estimated the breeding values based on pedigree data that does not have the power to recognize or select superior genomic segments. In this period, the superior genomic segments accumulated and segregated in the domestic population without being immediately recognized. The implementation of genomic selection in year 40 allowed for these segments to be recognized, selected, and promoted. This increased the annual genetic gain compared to when the genomic selection was not preceded by conventional selection (Figure S1). An additional boost also came from domestic reference population already containing foreign genomic segments. The described phenomena also changed the effect of varying the use of foreign sires. When both populations implemented genomic selection simultaneously, mating all domestic females with foreign sires yielded the highest annual genetic gain as well as the highest genetic gain in year 50 (Figure 1 and Figure S1). In contrast, when the populations ran a different scheme for ten years, using only foreign sires had the lowest annual genetic gain among the import scenarios (Figure S1). These scenarios used only foreign sires and thus did not benefit from accumulating superior genomic segments which later boosted the genetic gain of the genomic selection in other scenarios. Smith and Banos (1991) showed that when the initial genetic means of the populations differ, the population with the lower genetic mean can catch-up in three to five generations. We did not see this in our study. But, we saw that implementing domestic genomic selection with a 10-year delay allows for the annual domestic genetic gain in years 40-50 to exceed the annual foreign genetic gain (Figure S1).

### Implications

Cattle breeding programs import genetic material to increase genetic value, improve connectedness between countries, and increase genetic diversity. In this study, we focused on importing to increase genetic mean. We have simulated two genetically correlated populations where one is advanced and the other one is conservative. This is very common in dairy cattle breeding where most genetic improvement originates from only few populations and is disseminated by import (Gorjanc et al., 2012). When the impact of import is high and the breeding goals between populations are similar, the existence of local breeding programs must be reconsidered. The high cost of phenotyping and genotyping in the local population is difficult to justify when most of the genetic improvement is imported. Still, local multiplication of foreign genetic improvement can be economically reasonable and domestic phenotypes can be collected for the purpose of the management, essentially providing the required ingredients for a domestic breeding program.

When the correlation between the environments is high, import is beneficial for the genetic gain. This is the case in many dairy cattle breeding programs, but not for all (Interbull, 2021). For example, New Zealand’s Holstein population has a lower genetic correlation with other populations due to the grass-based system and a strict requirement for 365-day calving interval (Harris, 2005). Slagboom et al. (2019) showed that the break-even correlation with equally sized populations, above which it is more beneficial to have one joint instead of two separate breeding programs, is between 0.54 and 0.66. For populations of different size this correlation increases to 0.79 (Slagboom et al., 2019). However, differences in local environmental conditions can sometimes be overcome with an appropriate management that enables production with animals adjusted for other environments. This is the case for the Holstein population in Israel, which is genetically highly correlated to the Holstein population in other environments despite the differences between the actual environmental conditions. In Israel, breeding of dairy cattle on big farms was enabled by development of new technologies and management practices which are necessary for high production and reproduction (Flamenbaum and Galon, 2010). Still, there are many countries where these adjustments are more difficult to implement. For example, in India there is 16.5% of the world dairy cattle population with an average size between 0.6 and 2.0 cows per holding depending on the holding size (Food and Agriculture Organization of the United Nations, 2012; Landes et al., 2017). Small farms and severe climatic conditions do not enable establishment of the microenvironment necessary for a modern dairy cattle breeding. Furthermore, India is not part of routine Interbull international evaluation and genetic correlations of Indian dairy cattle population with the other countries are unknown.

In addition to importing, populations can increase genetic gain by changing their selection strategy, such as increasing the intensity of selection, or shortening the generation interval. Populations can also alleviate the loss of genetic variability and increase sustainability by implementing optimum contribution selection. In our previous study we tested the effect of varying the use of sires in a population equal to the one simulated here (Obšteter et al., 2019). We showed that small populations can increase the genetic gain of genomic selection by 25% by rapidly turning over elite sires. We also showed that the use of optimum contribution selection increases the sustainability of genomic selection and the efficiency of converting genetic variability to genetic gain. Further on, Fetherstone et al. (2021) showed that the genetic and economic benefits of importing foreign rams into the domestic sheep population were inferior to benefits achieved by increasing the use of progressive domestic selection.

Another strategy to increase genetic gain would be also to allow for selection across populations and exchange of information. In our population we assumed within population selection and a fixed extent of the foreign sire use in the domestic population. Small populations would likely benefit more by considering both domestic and foreign sires as potential sires and performing across population selection since this would increase selection intensity. This would also allow for a more flexible design in which the use of foreign sires would be optimized to maximize the genetic gain or other selection goals. However, selection across populations requires adjusting the evaluation methods to account for GxE, difference in selection goals, or even difference in breeds. Some of these aspects are adressed in so called multiple across-country evaluation (MACE) that is a national evaluation that integrates multi-national information (Schaeffer, 1994; Vandenplas et al., 2017). Furthermore, in genomic selection, collaboration between breeds can benefit from exchange of genomic information, either genomic data on selection candidates to increase reference population, or genotype-phenotype association. The latter is also valuable even when no animals are selected across environments (Andonov et al., 2017; Slagboom et al., 2019).

## CONCLUSIONS

In this study we explored the effect of import to genetic gain, genetic variability, and contributions of domestic and foreign selection to the genetic gain in a small cattle population. We showed that although using foreign sires can increase domestic genetic gain and genetic variability, increasing their use has diminishing returns. This means that there is an optimal point above which its better to use resources to support the domestic breeding actions. This point was between 25% and 50% in our simulation study. Above this point import did not increase genetic gain while it significantly increased the percentage of foreign genes in the domestic population. Such overreliance on import can decreases the importance of domestic breeding programs and the adaptation of animals to the local environmental conditions.

## ACKNOWLEDGEMENTS

JO acknowledges support from the ARRS Research program P4-0133. GG and IP acknowledge support from the BBSRC ISP grant BBS/E/D/30002275 to The Roslin Institute. JJ acknowledges the support from the Norwegian Research council through the project 309611. For the purpose of open access, the authors have applied a Creative Commons Attribution (CCBY) license to any Author Accepted Manuscript version arising from this submission.

## APPENDIX

**Table S1:** Pairwise p-values for differences in genetic gain in year 50 of scenarios where both populations implemented genomic selection simultaneously. The values below and above the diagonal compare the scenarios when genetic correlation between the populations was 0.8 and 0.9, respectively. Values on the diagonal display the genetic gain at genetic correlation 0.8 and 0.9, separated with a comma, and a p-value for the difference between the two values. Values in bold indicate statistically significant differences.

| Import | 0% | 10% | 25% | 50% | 100% | 100%_BD | ForeignPop<br>domestic | ForeignPop<br>foreign |
| --- | --- | --- | --- | --- | --- | --- | --- | --- |
| 0% | 4.56, 4.64<br>(0.50) | 1.00 | **0.04** | **<.0001** | **<.0001** | **0.03** | **<.0001** | **<.0001** |
| 10% | 1.00 | 4.63, 4.73<br>(0.47) | **0.04** | **<.0001** | **<.0001** | **0.03** | **<.0001** | **<.0001** |
| 25% | 0.99 | 1.00 | 4.66 5.12<br>(**0.00**) | 0.33 | 0.06 | 1.00 | 0.01 | **<.0001** |
| 50% | 0.96 | 1.00 | 1.00 | 4.69, 5.40<br>(**<.0001**) | 1.00 | 0.41 | 1.00 | **<.0001** |
| 100% | 1.00 | 1.00 | 1.00 | 1.00 | 4.55, 5.50<br>(**<.0001**) | 0.09 | 1.00 | **<.0001** |
| 100%_BD | 1.00 | 1.00 | 1.00 | 1.00 | 1.00 | 4.65, 5.14<br>(**0.00**) | **0.02** | **<.0001** |
| ForeignPop<br>domesticTrait | **0.00** | **0.00** | **0.00** | **0.00** | **0.00** | **0.00** | 4.53, 5.47<br>(**<.0001**) | **<.0001** |
| ForeignPop<br>foreignTrait | **<.0001** | **<.0001** | **<.0001** | **<.00010** | **<.0001** | **<.0001** | **<.0001** | 6.66, 6.36<br>(**<.0001**) |

**Table S2:** Pairwise p-values for differences in genic standard deviation in year 50, domestic population implemented genomic selection in year 40. The values below and above the diagonal compare the scenarios when genetic correlation between the populations was 0.8 and 0.9, respectively. Values on the diagonal compare the same scenario at two different genetic correlations between the populations. Values in bold indicate statistically significant differences.

| Import | 0% | 10% | 25% | 50% | 100% | 100%_BD | ForeignPop domestic | ForeignPop foreign |
| --- | --- | --- | --- | --- | --- | --- | --- | --- |
| 0% | 0.97, 0.96<br>(0.20) | 0.00 | <.0001 | **0.01** | <.0001 | <.0001 | <.0001 | <.0001 |
| 10% | 0.31 | 0.97, 0.98<br>(0.26) | 0.48 | 0.98 | 0.95 | 0.21 | 0.58 | 0.56 |
| 25% | **0.00** | 0.33 | 0.98, 0.98<br>(0.37) | 0.97 | **0.04** | 1.00 | 1.00 | 1.00 |
| 50% | <.0001 | 0.09 | 1.00 | 0.98, 0.98<br>(0.45) | 0.41 | 0.80 | 1.00 | 1.00 |
| 100% | 0.09 | 1.00 | 0.70 | 0.31 | 0.97, 0.97<br>(0.57) | **0.01** | **0.02** | **0.02** |
| 100%_BD | <.0001 | **0.00** | 0.47 | 0.84 | **0.01** | 0.99, 0.98<br>(0.51) | 0.88 | 0.87 |
| ForeignPop domesticTrait | <.0001 | 0.08 | 1.00 | 0.99 | 0.40 | 0.11 | 0.98, 0.98<br>(0.33) | 1.00 |
| ForeignPop foreignTrait | <.0001 | **0.12** | **1.00** | **0.97** | **0.50** | **0.08** | **1.00** | 0.98, 0.98<br>(<.0001) |

**Table S3:**
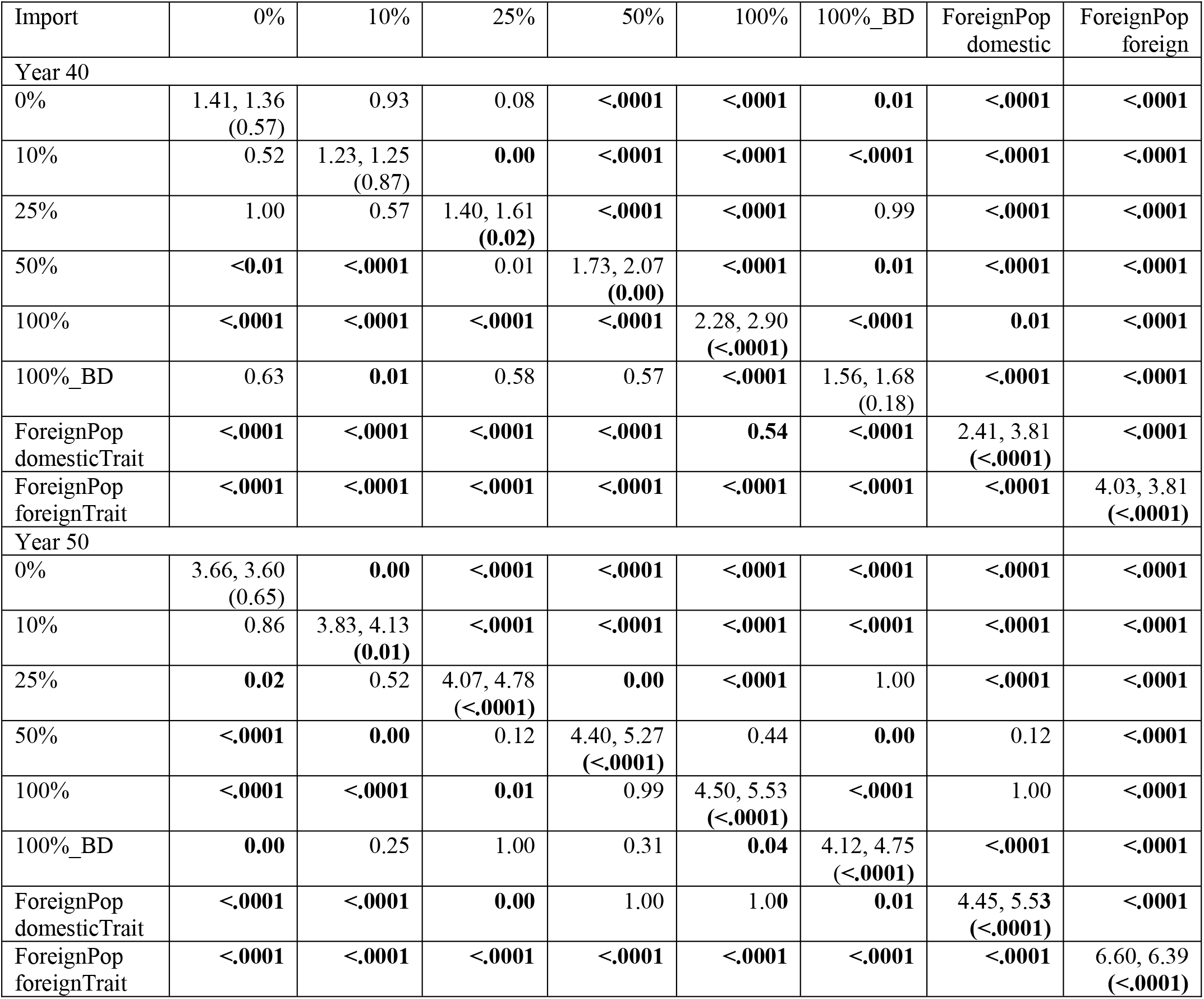
Pairwise p-values for differences in genetic gain in years 40 and 50, small population implemented genomic selection in years 40 The values below and above the diagonal compare the scenarios when genetic correlation between the populations was 0.8 and 0.9, respectively. Values on the diagonal compare the same scenario at two different genetic correlations between the populations. Values in bold indicate statistically significant differences.

| Import | 0% | 10% | 25% | 50% | 100% | 100%_BD | ForeignPop domestic | ForeignPop foreign |
| --- | --- | --- | --- | --- | --- | --- | --- | --- |
| Year 40 |  |  |  |  |  |  |  |  |
| 0% | 1.41, 1.36<br>(0.57) | 0.93 | 0.08 | <.0001 | <.0001 | 0.01 | <.0001 | <.0001 |
| 10% | 0.52 | 1.23, 1.25<br>(0.87) | 0.00 | <.0001 | <.0001 | <.0001 | <.0001 | <.0001 |
| 25% | 1.00 | 0.57 | 1.40, 1.61<br>(0.02) | <.0001 | <.0001 | 0.99 | <.0001 | <.0001 |
| 50% | <0.01 | <.0001 | 0.01 | 1.73, 2.07<br>(0.00) | <.0001 | 0.01 | <.0001 | <.0001 |
| 100% | <.0001 | <.0001 | <.0001 | <.0001 | 2.28, 2.90<br>(<.0001) | <.0001 | 0.01 | <.0001 |
| 100%_BD | 0.63 | 0.01 | 0.58 | 0.57 | <.0001 | 1.56, 1.68<br>(0.18) | <.0001 | <.0001 |
| ForeignPop domesticTrait | <.0001 | <.0001 | <.0001 | <.0001 | 0.54 | <.0001 | 2.41, 3.81<br>(<.0001) | <.0001 |
| ForeignPop foreignTrait | <.0001 | <.0001 | <.0001 | <.0001 | <.0001 | <.0001 | <.0001 | 4.03, 3.81<br>(<.0001) |
| Year 50 |  |  |  |  |  |  |  |  |
| 0% | 3.66, 3.60<br>(0.65) | 0.00 | <.0001 | <.0001 | <.0001 | <.0001 | <.0001 | <.0001 |
| 10% | 0.86 | 3.83, 4.13<br>(0.01) | <.0001 | <.0001 | <.0001 | <.0001 | <.0001 | <.0001 |
| 25% | 0.02 | 0.52 | 4.07, 4.78<br>(<.0001) | 0.00 | <.0001 | 1.00 | <.0001 | <.0001 |
| 50% | <.0001 | 0.00 | 0.12 | 4.40, 5.27<br>(<.0001) | 0.44 | 0.00 | 0.12 | <.0001 |
| 100% | <.0001 | <.0001 | 0.01 | 0.99 | 4.50, 5.53<br>(<.0001) | <.0001 | 1.00 | <.0001 |
| 100%_BD | 0.00 | 0.25 | 1.00 | 0.31 | 0.04 | 4.12, 4.75<br>(<.0001) | <.0001 | <.0001 |
| ForeignPop domesticTrait | <.0001 | <.0001 | 0.00 | 1.00 | 1.00 | 0.01 | 4.45, 5.53<br>(<.0001) | <.0001 |
| ForeignPop foreignTrait | <.0001 | <.0001 | <.0001 | <.0001 | <.0001 | <.0001 | <.0001 | 6.60, 6.39<br>(<.0001) |

**Table S4:** Pairwise p-values for differences in genic standard deviation in years 40 and 50, small population implemented genomic selection in year 40.The values below and above the diagonal compare the scenarios when genetic correlation between the populations was 0.8 and 0.9, respectively. Values on the diagonal compare the same scenario at two different genetic correlations between the populations. Values in bold indicate statistically significant differences.

| Import | 0% | 10% | 25% | 50% | 100% | 100%_BD | ForeignPop domestic | ForeignPop foreign |
| --- | --- | --- | --- | --- | --- | --- | --- | --- |
| Year 40 |  |  |  |  |  |  |  |  |
| 0% | 0.99, 0.99<br>(0.26) | 0.93 | 0.27 | 0.06 | 1.00 | **0.01** | **0.02** | **0.02** |
| 10% | 0.91 | 0.99, 0.99<br>(0.29) | 0.94 | 0.60 | 0.97 | 0.28 | 0.43 | 0.62 |
| 25% | 0.14 | 0.86 | 0.99, 0.99<br>(0.41) | 1.00 | 0.36 | 0.94 | 1.00 | 1.00 |
| 50% | **0.00** | 0.14 | 0.91 | 1.00, 1.00<br>(0.82) | 0.09 | 1.00 | 1.00 | 1.00 |
| 100% | 0.34 | 0.98 | 1.00 | 0.68 | 0.99, 0.99<br>(0.36) | **0.02** | **0.02** | **0.04** |
| 100%_BD | **<.0001** | 0.201 | 0.47 | 0.99 | 0.20 | 1.00, 1.00<br>(0.95) | 0.98 | 0.92 |
| ForeignPop domesticTrait | **0.00** | 0.06 | 0.95 | 1.00 | 0.67 | 0.78 | 0.99, 1.00<br>(0.35) | 1.00 |
| ForeignPop foreignTrait | **0.00** | 0.06 | 0.94 | 1.00 | 0.65 | 0.79 | 1.00 | 0.99, 1.00<br>(0.71) |
| Year 50 |  |  |  |  |  |  |  |  |
| 0% | 0.97, 0.96<br>(0.22) | **0.00** | **<.0001** | **0.04** | 1.00 | **<.0001** | **0.02** | **0.03** |
| 10% | **<.0001** | 0.98, 0.98<br>(0.79) | 1.00 | 0.90 | **0.01** | 0.98 | 0.26 | 0.18 |
| 25% | **<.0001** | 0.89 | 0.99, 0.99<br>(0.52) | 0.72 | **0.0** | 1.00 | 0.09 | 0.06 |
| 50% | **0.00** | 1.00 | 0.72 | 0.98, 0.98<br>(0.83) | 0.24 | 0.31 | 1.00 | 0.99 |
| 100% | 0.34 | 0.14 | **0.00** | 0.28 | 0.98, 0.98<br>(0.79) | **0.00** | 0.21 | 0.29 |
| 100%_BD | **<.0001** | 0.15 | 0.90 | 0.07 | **<.0001** | 0.99, 0.99<br>(0.99) | **0.01** | **0.00** |
| ForeignPop domesticTrait | **<.0001** | 0.95 | 0.08 | 1.00 | 0.28 | **0.00** | 0.98, 0.98<br>(0.14) | 1.00 |
| ForeignPop foreignTrait | **<.0001** | 0.92 | 0.06 | 0.99 | 0.34 | **0.00** | 1.00 | 0.98, 0.98<br>(0.58) |

**Figure S1:**
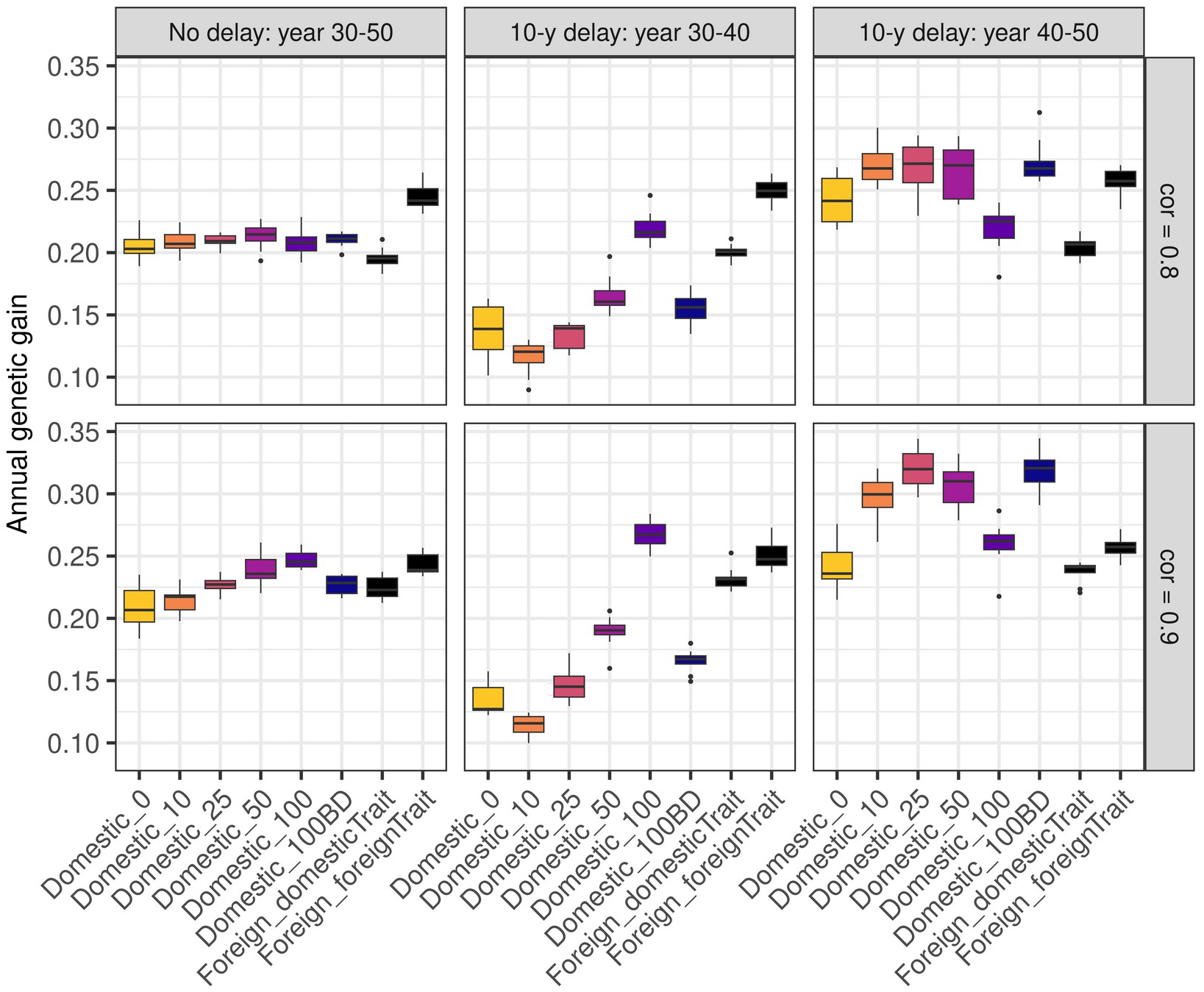
Annual genetic gain by scenario, genetic correlation between populations, and time of implementing genomic selection. The scenarios named “Domestic” show the domestic genetic gain in scenarios with varying percentage of females mated with foreign sires (0, 10, 25, 50, 100). Scenario 100BD mated only bull dams with foreign sires. The scenario named “Foreign” shows the genetic gain of the foreign population for the trait expressed on the scale of the domestic (“domesticTrait”) and the foreign (“foreignTrait”) breeding program. For the scenarios that implemented the genomic selection in the domestic population with a 10-year delay, we show the annual genetic gain of the years 30-40 (conventional selection in the domestic population) and years 40-50 (genomic selection in the domestic population) separately.

**Figure S2:**
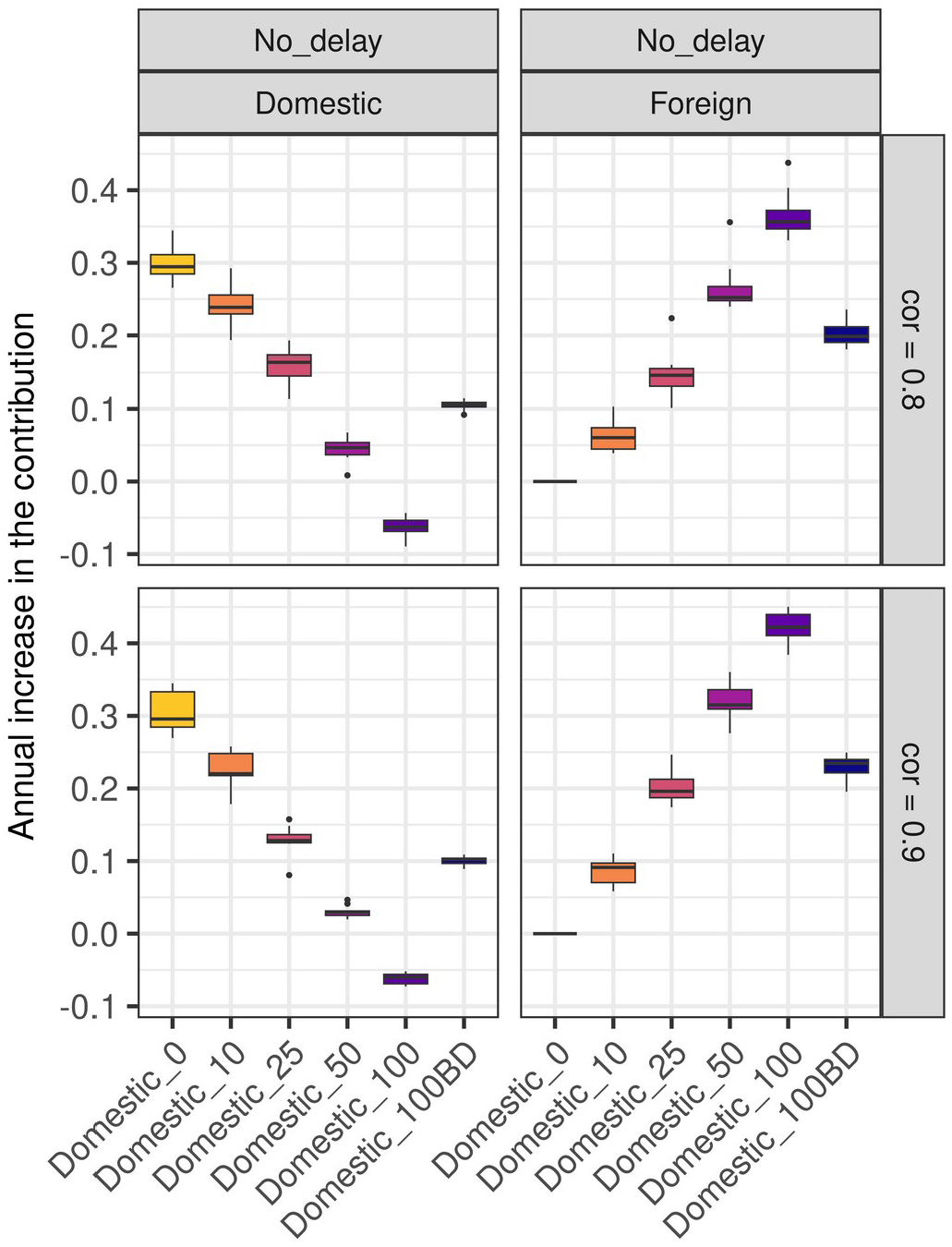
Average annual rate in the contribution of domestic and foreign selection in scenarios that simultaneously implemented genomic selection in both populations. The scenarios named “Domestic” show the domestic genetic gain in scenarios with varying percentage of females mated with foreign sires (0, 10, 25, 50, 100). Scenario 100BD mated only bull dams with foreign sires. The scenario named “Foreign” shows the genetic gain of the foreign population for the trait expressed on the scale of the domestic (“domesticTrait”) and the foreign (“foreignTrait”) breeding program.

**Figure S3:**
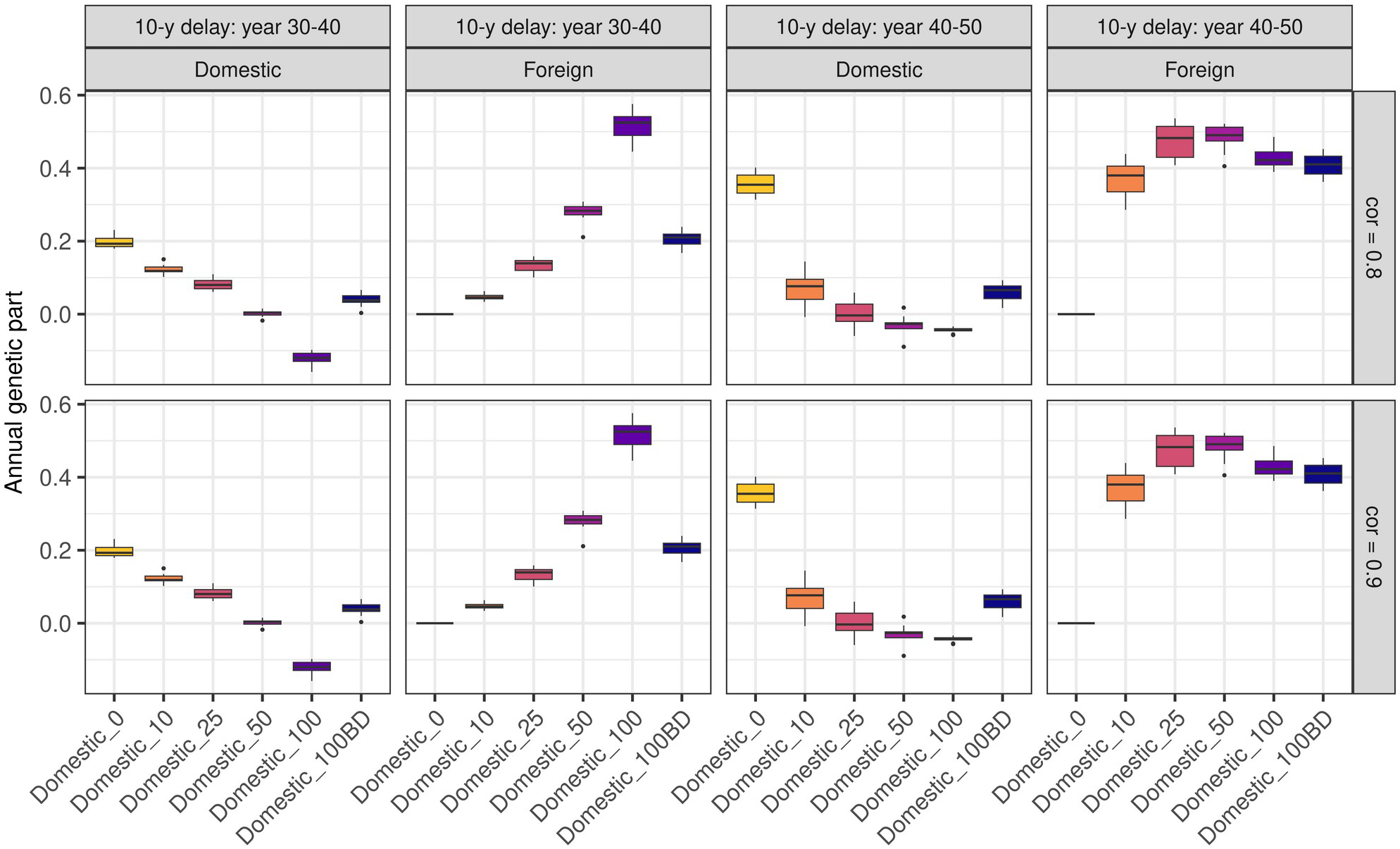
Annual change in the contribution of domestic and foreign selection in scenarios that implemented genomic selection in the domestic population 10 years after the foreign population. The scenarios named “Domestic” show the domestic genetic gain in scenarios with varying percentage of females mated with foreign sires (0, 10, 25, 50, 100). Scenario 100BD mated only bull dams with foreign sires. The scenario named “Foreign” shows the genetic gain of the foreign population for the trait expressed on the scale of the domestic (“domesticTrait”) and the foreign (“foreignTrait”) breeding program.

**Figure S4:**
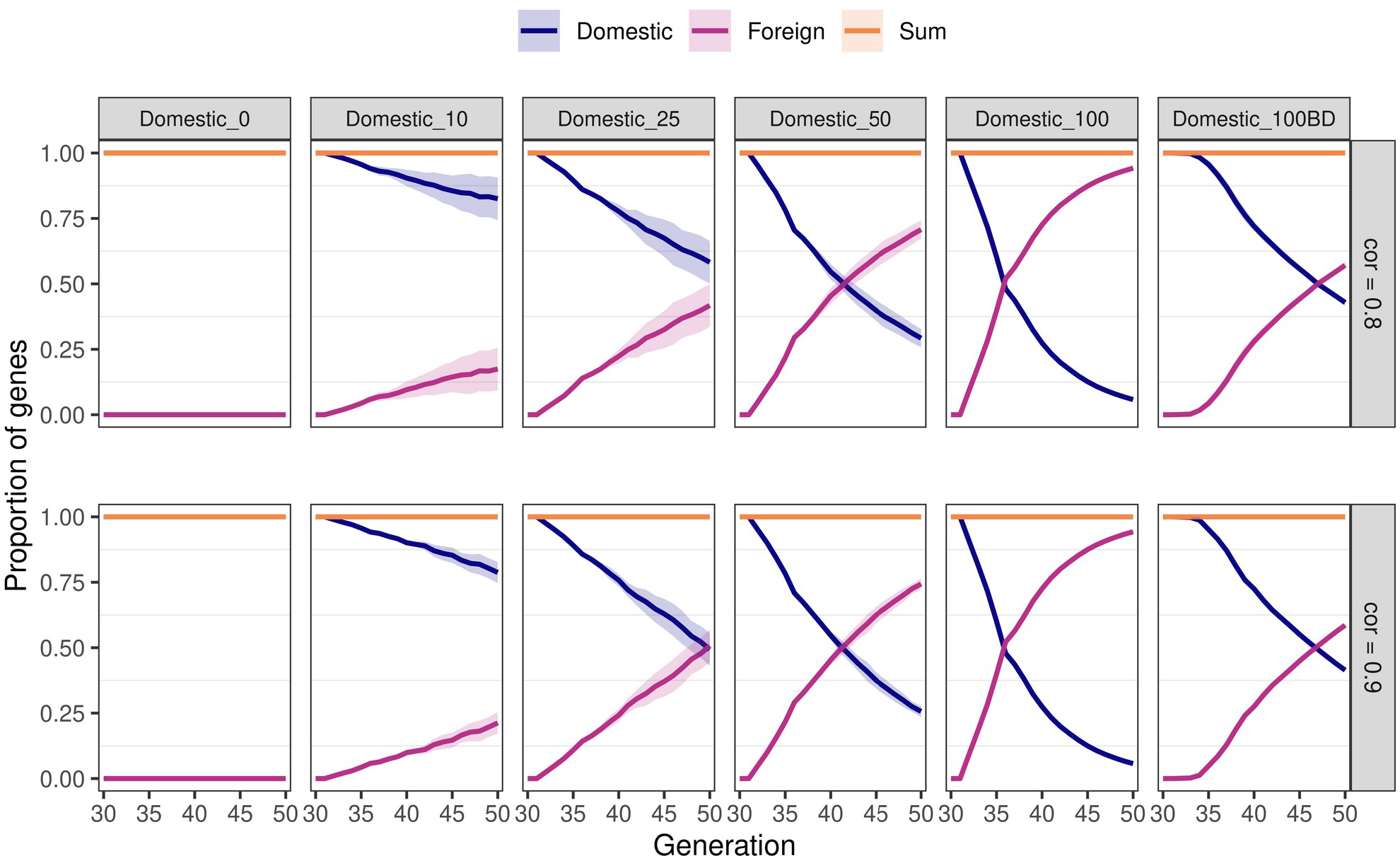
Proportion of domestic and foreign genes in the domestic population in scenarios with simultaneous implementation of genomic selection in both populations. The scenarios first varied the percentage of females mated with foreign sires (0, 10, 25, 50, 100). Scenario 100BD mated only bull dams with foreign sires. Second, the scenarios varied genetic correlation between two breeding programs (0.8 or 0.9).

**Figure S5:**
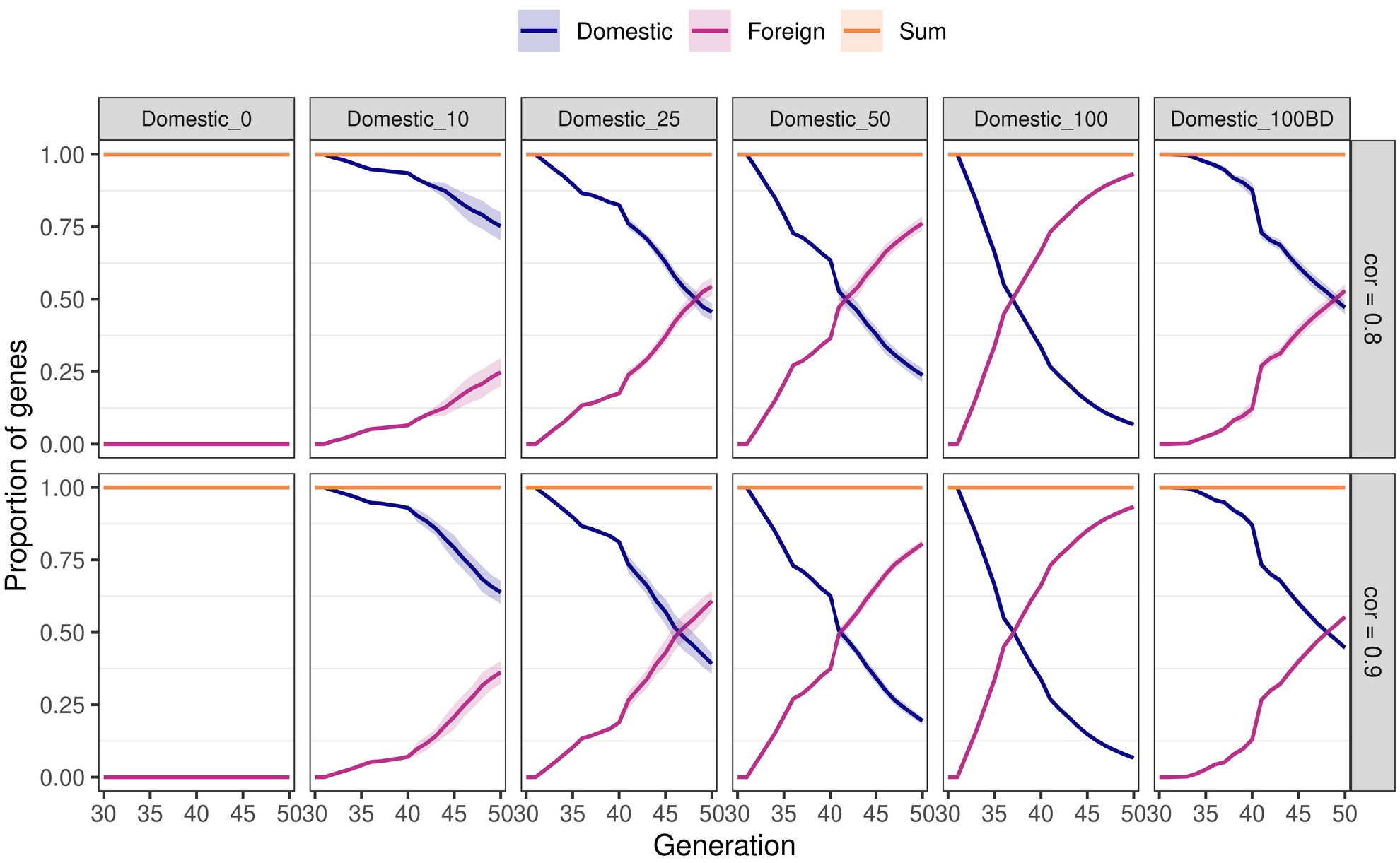
Proportion of domestic and foreign genes in the domestic population in scenarios with a 10-year delay in the implementation of genomic selection in the domestic population. The scenarios first varied the percentage of females mated with foreign sires (0, 10, 25, 50, 100). Scenario 100BD mated only bull dams with foreign sires. Second, the scenarios varied genetic correlation between two breeding programs (0.8 or 0.9).

**Figure S6:**
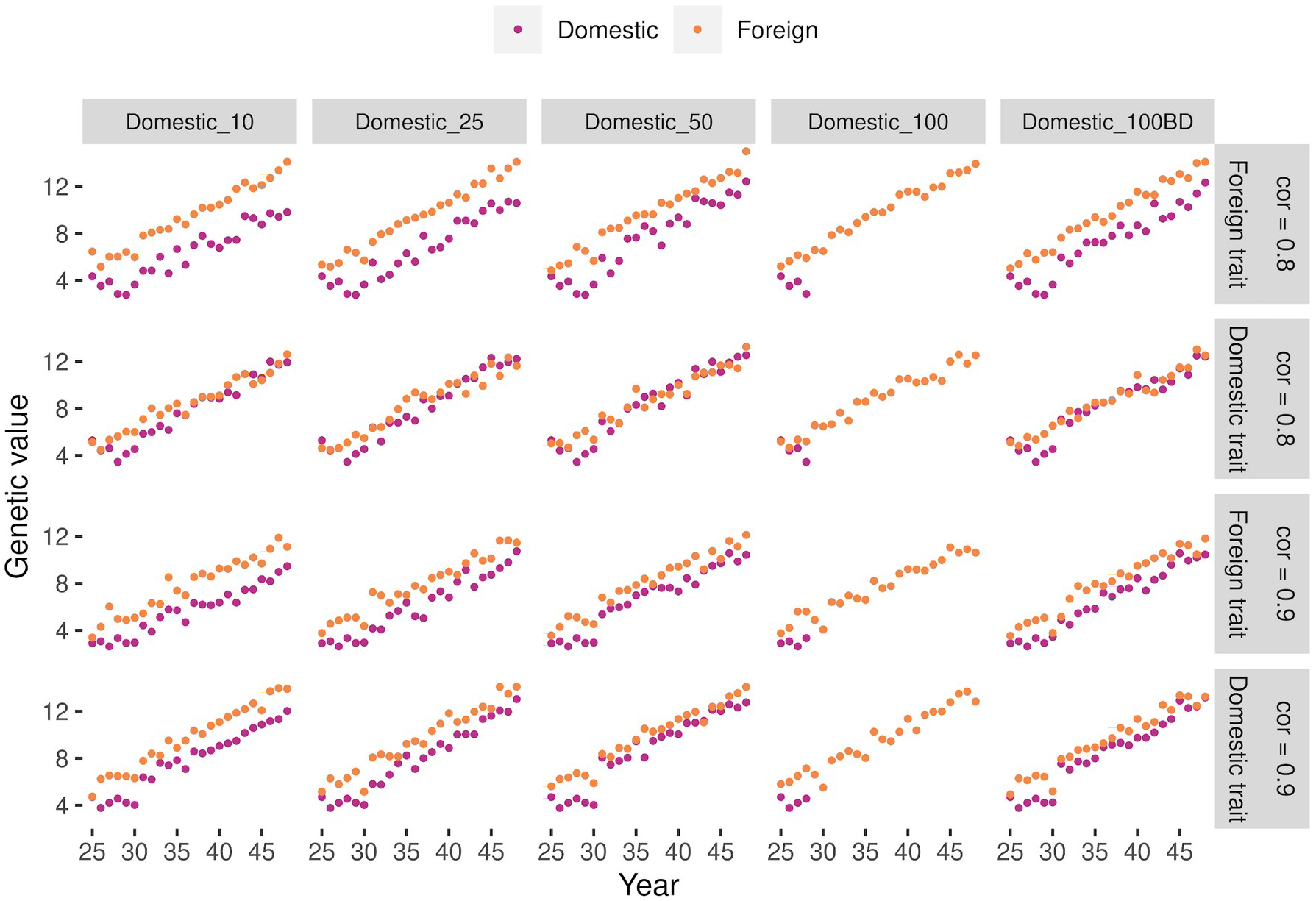
Genetic values of foreign and domestic sires used in the domestic population for the trait expressed in the domestic and foreign population for one simulation replicate. Figure shows scenarios with simultaneous implementation of genomic selection in both populations. The scenarios first varied the percentage of females mated with foreign sires (0, 10, 25, 50, 100). In 100BD scenario we mated only bull dams with foreign sires. Second, the scenarios varied genetic correlation between two breeding programs (0.8 or 0.9).

## REFERENCES

Andonov S., D. A. L. Lourenco, B.O. Fragomeni, Y. Masuda, I. Pocrnic, S. Tsuruta, I. Misztal. 2017. Accuracy of breeding values in small genotyped populations using different sources of external information—A simulation study. J Dairy Sci, 100, 1:395–401, https://doi.org/10.3168/jds.2016-11335

Banos G., C. Smith. 1991. Selecting bulls across countries to maximize genetic improvement in dairy cattle. J Anim Breed Genet, 108, 1-6: 174–181, https://doi.org/10.1111/j.1439-0388.1991.tb00172.x

Cao L., H. Liu, H. A. Mulder, M. Henryon, J. R. Thomasen, M. Kargo, A.C. Sørensen. 2020. Genomic breeding programs realize larger benefits by cooperation in the presence of genotype × environment interaction than conventional breeding programs. Front Genet, 11: 251, https://doi.org/10.3389/fgene.2020.00251

Falconer D.S., T. F. C. Mackay. 1996. Introduction to quantitative genetics. 4th ed.

Harlow, Longman Fikse W. F., R. Rekaya, K. A. Weigel. 2003. Genotype×environment interaction for milk production in Guernsey cattle. J Dairy Sci, 86, 5: 1821–1827, https://doi.org/10.3168/jds.S0022-0302(03)73768-0

Flamenbaum I., N. Galon. 2010. Management of heat stress to improve fertility in dairy Cows in Israel. J Reprod Dev, 56, Suppl: S36–S41, https://doi.org/10.1262/jrd.1056S36

Food and Agriculture Organization of the United Nations. 2012. FAOSTAT statistical database. http://www.fao.org/faostat/en/#data

Gorjanc G., F. Hely, P. Amer. 2012. Partitioning international genetic trends by origin in Holstein bulls. ICAR. https://www.icar.org/index.php/icar-meetings-news/38th-session-cork-ireland/

Gorjanc G., K. Potočnik, L. A. García-Cortés, J. Jakobsen, J. Dürr. 2011. Partitioning of international genetic trends by origin in Brown Swiss bulls. V: Proceedings of the 2011 Annual Interbull Open Meeting – Interbull Bulletin 44. Interbull Meeting, Stavanger, 27th – 29th August 2011. Hossein Jorjani (ur.). Uppsala, Interbull Centre: 272 p.

Harris B. L. Breeding dairy cows for the future in New Zealand. 2005. N Z Vet J, 53, 6: 384–389, https://doi.org/10.1080/00480169.2005.36582

Interbull. 2021. MACE evaluations archive. https://interbull.org/ib/maceev_archive (17th February 2021)

Landes M., J. Cessna, L. Kuberka, K. Jones. 2017. India’s dairy sector: structure, performance, and prospects, LDPM-272-01. United States Department of Agriculture, Economic Research Service, https://www.ers.usda.gov/webdocs/outlooks/82639/ldpm-272-01.pdf?v=2655.9

Misztal I., S. Tsuruta, D. A. L. Lourenco, Z. Masuda, I. Aguilar, A. Legarra, Z. Vitezica. 2018. Manual for BLUPF90 family programs. University of Georgia: 142 str. http://nce.ads.uga.edu/wiki/doku.php?id=documentation

Mulder H. A., P. Bijma. 2006. Benefits of cooperation between breeding programs in the presence of genotype by environment interaction. J Dairy Sci, 89, 5: 1727–1739, https://doi.org/10.3168/jds.s0022-0302(06)72241-x

Mulder, H. A. 2016. Genomic Selection Improves Response to Selection in Resilience by Exploiting Genotype by Environment Interactions. Front Genet, 7, 178:1–11, https://doi.org/10.3389/fgene.2016.00178

Obšteter J., J. Jenko, J. M. Hickey, G. Gorjanc. 2019. Efficient use of genomic information for sustainable genetic improvement in small cattle populations. J Dairy Sci, 102, 11: 9971–9982, https://doi.org/10.3168/jds.2019-16853

Obšteter J., J. Holl, J. M. Hickey, G. Gorjanc. 2021. AlphaPart - R implementation of the method for partitioning genetic trends. Genet Sel Evol, 53, 30:1–11, https://doi.org/10.1186/s12711-021-00600-x

Ojango J. M. K., G. E. Pollott. 2002. The relationship between Holstein bull breeding values for milk yield derived in both the UK and Kenya. Livest Prod Sci, 74, 1: 1–12, https://doi.org/10.1016/S0301-6226(01)00282-2

Rekaya R., K. A. Weigel, D. Gianola. 2001. Application of a structural model for genetic covariances in international dairy sire evaluations. J Dairy Sci, 84, 6: 1525–1530, https://doi.org/10.3168/jds.S0022-0302(01)70186-5

Robertson A. 1959. The sampling variance of the genetic correlation coefficient. Biometrics, 15, 3: 469–485, https://doi.org/10.2307/2527750

Schaeffer L. R. 1994. Multiple-country comparison of Dairy Sires. J Dairy Sci, 77, 9:2671–2678, https://doi.org/10.3168/jds.S0022-0302(94)77209-X

Schmidtmann C., M. Slagboom, A. C. Sørensen, D. Hinrichs, G. Thaller, M. Kargo. 2022..Short- and long-term consequences of collaboration between Northern European Red dairy and dual-purpose cattle. J Anim Breed Genet, 139, 4:447–461, https://doi.org/10.1111/jbg.12672

Smith C., G. Banos. 1991. Selection within and across populations in livestock improvement. J Anim Sci, 69, 6: 2387–2394, https://doi.org/10.2527/1991.6962387x

Slagboom M., M. Kargo, A.C. Sørensen, J. R. Thomasen, H. A. Mulder. 2019. Genomic selection improves the possibility of applying multiple breeding programs in different environments. J Dairy Sci, 102, 9: 8197–8209, https://doi.org/10.3168/jds.2018-15939

Slagboom M., A. C. Sørensen, J. R. Thomasen, H. Liu, M. Kargo, L. Hjortø. 2021. Ignoring genotype by environment interaction in the genetic evaluation of dairy cattle reduces accuracy but may increase selection intensity. J Dairy Sci, 104, 12:12756–12764. https://doi.org/10.3168/jds.2021-20876

Vandenplas J., M. Spehar, K. Potocnik, N. Gengler, G. Gorjanc. 2017. National single-step method that intergrates multi-national genomic information. J Dairy Sci, 100, 1:465–478, https://doi.org/10.3168/jds.2016-11733

Vandenplas J., M. P. L. Calus, G. Gorjanc. 2018. Genomic Prediction Using Individual-Level Data and Summary Statistics from Multiple Populations. Genetics, 210, 1: 53–69. https://doi.org/10.1534/genetics.118.301109

Wiggans G. R., J. B. Cole, S. M. Hubbard, T. S. Sonstegard. 2017. Genomic selection in dairy cattle: The USDA experience. Annu Rev Anim Biosci, 8, 5:309–327, https://doi.org/10.1146/annurev-animal-021815-111422

